# Mat-aCGH: a Matlab toolbox for simultaneous multisample aCGH data analysis and visualization

**DOI:** 10.1101/028761

**Authors:** Majid Mohammadi, Hossein Sharifi Noghabi

## Abstract

Mat-aCGH is an application toolbox for analysis and visualization of microarray-comparative genomic hybridization (array-CGH or aCGH) data which is based on Matlab. Full process of aCGH analysis, from denoising of the raw data to the visualization of the desired results, can be obtained via Mat-aCGH straightforwardly. The main advantage of this toolbox is that it is collection of recent well-known statistical and information theoretic methods and algorithms for analyzing aCGH data. More importantly, the proposed toolbox is developed for multisample analysis which is one of the current challenges in this area. Mat-aCGH is convenient to apply for any format of data, robust against diverse noise and provides the users with valuable information in the form of diagrams and metrics. Therefore, it eliminates the needs of another software or package for multisample aCGH analysis. aCGH Matlab source codes and datasets are freely available and can be downloaded at: *hsharifi.student.um.ac.ir/imagesm/*14407*/Mat–aCGH.rar*.

## 1. Introduction

We are in the post-genome era which means urgent need for computational methods for analyzing high-throughput genomics data in order to extract biological knowledge out of them (Zhou et al. 14). High-throughput technologies such as Next Generation Sequencing (NGS) [5] have paved the way for scientists to understand complex diseases in much better ways. One of these novel technologies is accentuation of comparative genomic hybridization (CGH) and microarray in arrayCGH (aCGH) [8]. aCGH is a high-throughput and high-resolution approach for measuring changes of copy numbers in thousands of DNA regions [6].

Investigation of DNA copy number aberrations in different diseases such as cancer has been truly enhanced via aCGH [7]. The primary goal of analyzing aCGH data is to identify the sections of copy number variation (CNV) and moreover, quantifying their amount at thousands of locations in a genome [11]. Although lots of studies have performed so far on aCGH analysis, most of them were based on single-sample [6]. Today, the focus of aCGH research is on multi-sample analysis [10], therefore, it is vital and crucial to provide researchers with a reliable toolbox in this regard. Previous attempts such as ArrayCyGHt [2] or CGHAnalyzer [3] provides the users with effective techniques, however, it is essential to propose novel toolboxes that take advantage of more recent methods with especial focus on robust denoising capabilities and multi-sample analysis.

## 2. Streamlined Steps

### 2.1. Loading Data

The methods provided in this toolbox can analyze any data loadable in Matlab. Aside from that, simulated data can be generated with adjustable number of dimensions and samples. To better distniguish the performance of various methods, the possiblity to contaminate the simulated data is considered and various types of noise are accomodated in this toolbox. Further, two types of aberrations are considered: the aberrations for each individual profile and aberrations shared among all profiles. The ratio between shared aberrations and all aberrations is called *shared percentage* (Zhou et al. 12). It is worth mentioning that any individual aCGH sample lies on the columns of the data matrix.

### 2.2. Analysis

After loading CGH array data or generating simulated profiles, there are five methods by which real or simulated data can be analyzed: TVSp (Zhou et al. 12), RPLA (Zhou et al. 13), Group Fused Lasso segmentation (GFLseg) (Bleakley and Vert 1), SHQ and LRHQ (Mohammadi and Abed Hodtani 4). TVSp approximated the whole aCGH profiles by minimizing nuclear norm of the whole matrix and the total variation of each sample. With the same approach, RPLA tried to recover true aCGH data by imposing an sparse error assumption on each sample. This method is claimed to be more robust in dealing with diverse noise. In contrast to TVSp and RPLA, GFLseg denoised aCGH data by minimizing the total variation of the entire matrix. In order to deal with different types of noise (especially non-Gaussian noise), SHQ and LRHQ accomodated different M-estimation loss functions. SHQ encouraged the approximated aCGH profiles to be as sparse and piecewise constant as possible and LRHQ proposed to recovere a low rank and piecewise constant aCGH data matrix.

### 2.3. Visualization

In order to visualize this toolbox, we provide the users with numerous informative plots and diagrams. In the case of the real example profiles, plots including: heat diagram, bar diagram, real (noisy) and recovered signals are provided. In addition to these plots and diagrams, different metrics such as receiver operating characteristic (ROC), accuracy and false discovery rate (FDR) are also included in Mat-aCGH. It is important to note that the latter metrics are only applicable for simulated examples and inputs. Fig. 1 illustrates aforementioned visualization for both simulated and real data. Fig. 1 (a) shows the ROC on a generated data with exponential noise. The different SNRs and shared percentage are considered and the results of recovered profiles are shown. Fig. 1 (b) dedicated to diagrams for Pollack dataset [9]. The first row plots heat diagram, the second row shows an individual sample of dataset before and after processing by LRHQ and the last row illustrates the bar diagram of the mentioned dataset.

**Figure 1:**
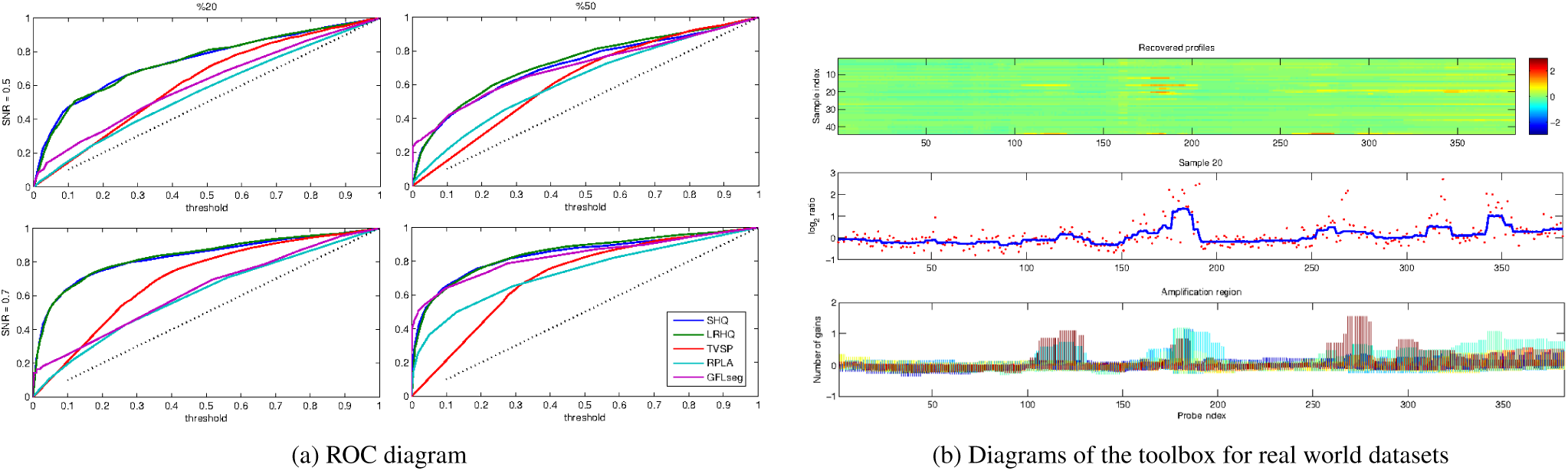
Toolbox visualization diagrams: (a) ROC diagram on simulated data contaminated with Exponential noise and different SNRs and shared percentage; (b) Bar and heat diagrams of Pollack et al. dataset [9] and one of its real and recovered samples; Red dots: real profile; Blue line: recovered profile

## 3. Conclusion

Although the provided toolbox is not covering all of the state-of-the-art methods and algorithms, it tried to make the best use of the best methods and deals with aCGH data analysis from different aspects and prospective. This Matlab-based toolbox can play a pivotal role in aCGH profiles denoising, analysis and visualization. To our knowledge, this is the first multisample aCGH toolbox that brought most recent methods and algorithms from information theoretic learning (ITL) to statistical learning. Mat-aCGH is completely user-friendly and any users with basic understanding of Matlab can apply it for wide range of data types readable in Matlab. In this version of Mat-aCGH, the users can load simulated and real examples provided by the authors or load their own dataset and compare the results with all of the methods implemented in this toolbox at the same time. For future, we are planning to develop Mat-aCGH in C and R and provide the users with graphical user interface.

